# Differences in functional connectivity distribution after transcranial direct-current stimulation: a connectivity density point of view

**DOI:** 10.1101/2020.11.23.395160

**Authors:** Bohao Tang, Yi Zhao, Archana Venkataraman, Kyrana Tsapkini, Martin A Lindquist, James Pekar, Brian Caffo

**Affiliations:** Maryland, United States

**Keywords:** Functional Connectivity, Density Regression, Random Graph

## Abstract

In this manuscript we consider the problem of relating functional connectivity measurements viewed as statistical distributions to outcomes. We demonstrate the utility of using the distribution of connectivity on a study of resting state functional magnetic resonance imaging association with an intervention. Specifically, we consider 47 primary progressive aphasia (PPA) patients with various levels of language abilities. These patients were randomly assigned to two treatment arms, tDCS (transcranial direct-current stimulation and language therapy) vs sham (language therapy only), in a clinical trial. We propose a novel approach to analyze the effect of direct stimulation on functional connectivity. We estimate the density of correlations among the regions of interest (ROIs) and study the difference in the density post-intervention between treatment arms. We discover that it is the tail of the density, rather than the mean or lower order moments of the distribution, that demonstrates a significant impact in the classification. This approach has several benefits. Among them, it drastically reduces the number of multiple comparisons compared to edge-wise analysis. In addition, it allows for the investigation of the impact of functional connectivity on the outcomes where the connectivity is not geometrically localized.

## 1. Introduction

The study of resting state brain connectivity via functional magnetic resonance imaging (fMRI) involves the investigation of correlations between cortical seeds, regions or voxels (henceforth referred to as foci). Friston, in particular, defined functional connectivity as the correlations, over time, between spatially distinct brain regions [1]. Nearly all inter-subject investigations of connectivity have focused on *localized correlations*. That is, they consider correlations between foci treated consistently across subjects. Mathematically, this can be described as saying that the methods are not invariant to subject-specific relabeling of the foci. In fact, for most methods, such as pairwise regressions on correlations across subjects or decomposition methods, shuffling foci labels within subjects is a form of null distribution. Furthermore, this lack of invariance applies regardless of the degree of granularity of the analysis (seed, region, voxel …) [1, 2, 3]. The methods and choice of granularity all center the focus on geographic consistency of correlations across groups of similar subjects. A notable exception is some variations of graph theory based methods, where graphical summaries may not be localized across subjects in the sense of being invariant to subject-specic foci labels [4, 5].

In this manuscript, we consider the distribution of resting state correlations and how these correlations vary between treatment arms. This form of density regression has several benefits. A primary one is the relaxation of the consistent localization assumption across subjects. Specifically, localization analyses makes the, often unchallenged, assumption that pairs of foci represent the same correlated functional specialization across exchangeable subjects. This assumption is grounded in the neurological theory of functional specialization dating back to the foundational works of Broca and Weirnicke [6, 7]. However, it is clear that in specific applications and biological settings, the neural geography of functional specialization can vary. As an extreme example, subjects with brain damage in their youth often have the neuroplasticity that remaps a function to atypical areas [8].

There are existing studies that focus on utilizing the distribution of resting state correlations. For example, Petersen [9] considers the distribution of correlations between a seed voxel and all other voxels within a region of interest (ROI), to summarise the state of such ROI. Also, Scheinost [10] further considered such distributions across all pairs of voxels. This work derived a degree function from the connection density as a summary of the connectivity of each voxel. As a result, the study continues to focus on localized effects, where the use of the connectivity density is mainly to achieve a more informative localized summary of brain connectivity.

Our study is motivated by a resting-state fMRI study of primary progressive aphasia (PPA) patients, where it is feasible to relax the geometric localization assumption.In the study, the patients were randomly assigned into two treatment groups, tDCS (transcranial direct-current stimulation [11] + language therapy) and sham (language therapy only). In the tDCS group, the stimulation target, the left inferior frontal gyrus (IFG), is less likely to satisfy the across-subject localization assumption due to spatial normalization and brain functional specialization. In addition, the stimulation electrode patches were big, 5 × 5 = 25 cm^2^, thus, the stimulation areas were extended beyond the left IFG. This may induces additional variation across subjects, and thus may result in violations of localization assumptions. Here, we propose a novel approach to represent the effect of stimulation on functional connectivity. By ignoring the spacial heterogeneity, we directly study the change on the distribution of correlation between the regions of interest (ROIs) and therefore the approach has the potential to be highly robust to spatial registration.

In following section 2, we will introduce the experimental design and our approach. Results both for simulation and real data will be shown in section 3. And section 4 contains an overall discussion about the paper.

## 2. Material and Methods

### 2.1. Experimental Design

The data analyzed in this study were part of a larger crossover study on aphasia treatment using tDCS. All of the analyzed subjects had at least two years of progressive language deficit and no history of any other neurological condition that may have affected their language ability. Subjects had atrophy predominantly in the left hemisphere. Subjects were diagnosed via neuropsychological testing, language testing, MRI and clinical assessment according to consensus criteria [12]. The study was approved by the Johns Hopkins Hospital Institutional review board and all subjects provided informed consent to participate in the study.

A total of 50 right handed, native English speaking patients had a pre-intervention scan (scan1), 48 had a post-intervention scan (scan2). One patient was deleted from the analysis because of missing values in the connectivity matrix. Among the remaining 47 post-intervention scanned patients, 25 had transcranial direct-current stimulation + language therapy and the remaining 22 patients had only language therapy. Several baseline covariates were recorded including: gender, disease onset (years), age at the start of therapy and language severity. These patients were diagnosed with three variant types, including the logopenic, the nonfluent, and the semantic, based on brain functions compromised, which reflects brain areas that show initial atrophy. Patients with *Logopenic* variant PPA (lvPPA) present with word-finding difficulties and disproportionately impaired sentence repetition. Patients with *nonfluent* variant PPA (nfvPPA) present with difficulty producing grammatical sentences and/or motor speech impairment (apraxia of speech). Finally, patients with *semantic* variant PPA (svPPA) present with fluent speech, but impaired word comprehension. See Table 1 for a summary of demographic and clinical information on the participants.

**Table 1:** Patient demographics. For age, years post onset, severity, values shown are mean (standard deviation). P-values are from the Welch two sample t-tests for continuous outcomes and Fisher’s exact test for categorical outcomes. Language severity is based on the language subset from the FTD-CDR scale. Total severity refers to the sum of boxes, including language and behavior as added in [13].

|  | Combined (n = 47) | tDCS (n = 25) | Sham (n = 22) | P-value |
| --- | --- | --- | --- | --- |
| Sex | 22F, 25M | 11F, 14M | 11F, 11M | 0.773 |
| PPA variant | 15L, 23N, 9S | 9L, 12N, 4S | 6L, 11N, 5S | 0.801 |
| Age | 67.3 (6.8) | 65.8 (8.1) | 69.1 (5.0) | 0.146 |
| Year post onset | 4.2 (2.8) | 4.3 (3.2) | 4.0 (2.3) | 0.722 |
| Language severity | 1.7 (0.8) | 1.7 (0.9) | 1.8 (0.8) | 0.719 |
| Total severity | 6.3 (4.5) | 5.7 (3.9) | 7.0 (5.2) | 0.597 |

### 2.2. Data Preprocessing

MRI scans were obtained at the Kennedy Krieger Institute at Johns Hopkins University, using a 3 T Philips Achieva MRI scanner equipped with a 32-channel head coil. Resting-state fMRI (rsfMRI) data were acquired for approximately 9 min (210 time-point acquisitions) post-intervention. We used a 2D EPI sequence with SENSE partial-parallel imaging acceleration to obtain an in-plane resolution of 3.3 × 3.3 mm^2^ (64 × 64 voxels; TR/TE = 2500/30 ms; flip angle = 75°; SENSE acceleration factor = 2; SPIR for fat suppression, 3 mm slice thickness). The data were co-registered with structural scans into the same anatomical space. Structural scans, acquired axially with a scan time of 6 min (150 slices), used a T1-weighted MPRAGE sequence with 3D inversion recovery, magnetization-prepared rapid gradient, isotropic with a resolution of 1 × 1 × 1 mm^3^ (FOV = 224 × 224 mm^2^; TR/TE = 8.1/3.7 ms; flip angle = 8°; SENSE acceleration factor = 2).

Using MRICloud, a cloud-platform for automated image parcellation approach (atlas-based analysis (ABA)), the MPRAGE scan was parcelled into 283 structures [14]. In detail, each participant’s high resolution MPRAGE was segmented by using a multi-atlas fusion label algorithm (MALF) and large deformation diffeomorphic metric mapping, LDDMM [15, 16, 17]. This highly accurate diffeomorphic algorithm, associated with multiple atlases, minimizes the mapping inaccuracies due to atrophy or local shape deformations. All analyses were performed in native space. To control for relative regional atrophy, volumes for each ROI were normalized by the total intracerebral volume (total brain tissue without myelencephalon and cerebrospinal fluid). The resting-state fMRI was also processed in MRICloud and analyzed in a seed-by-seed manner. The image processing was described in our previous publication [18] including routines imported from the SPM connectivity toolbox for coregistration, motion, and slice timing correction; physiological nuisance correction using CompCor [19]; and motion and intensity TR outlier rejection using “ART” (https://www.nitrc.org/projects/artifact_detect/). The MRICloud pipeline follows well established steps for rsfMRI processing: after exclusion of “outlier” TRs, detected by ART routine (parameters: 2 standard deviations for motion and 4 standard deviations for intensity, more severe than the default of 9), the movement matrix combined with the physiological nuisance matrix is used in the deconvolution regression for the remaining TRs. These two steps for motion correction (outlier rejection and regression of motion parameters) ensure the minimization of the motion effect. The parcels resultants from the high resolution T1 segmentation were brought to the resting state dynamics by co-registration. Time-courses of 78 cortical and deep gray matter ROIs were extracted and the correlations among them were calculated.

### 2.3. Density regression

We propose to quantify the effect of possibly non-localized stimulation on functional connectivity through a density regression. We make the assumption that the connectivity matrix **C**_*i*_ of patient *i* is the adjacency matrix of a random weighted graph, for *i* = 1*, …, n*, and *n* is the number of subjects. For all pairs of elements in the set {(*s, t*; *i*) | *s* < *t* ≤ *N*}, where *N* is the number of nodes in the graph, we have:

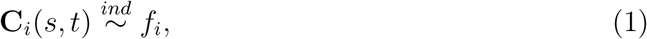

where *f_i_* is a density function. We refer to this density as the *connectivity density* of subject *i*. The process of proceeding from fMRI scans to the connectivity density is outlined in Figure 1. We estimate connectivity matrix from temporal correlation of BOLD signals between regions of interest (ROIs) after parcellation. And then estimate the connectivity density. In practice, one can use the vectorized elements in the upper triangular portion of the connectivity matrix to estimate the density using smoothing splines [20], which performs a maximum likelihood estimation on the spline coefficients for estimating the logarithm of the density function under a smoothness penalty. We choose this approach as it directly returns the splines, which are both mathematically and practically convenient, especially for performing a functional regression. In addition, it sets a boundary of the support for the estimated density, which is beneficial to our case as correlation coefficients are bounded between −1 and 1. Kernel density estimators [21] are also implemented as a comparison.

**Figure 1:**
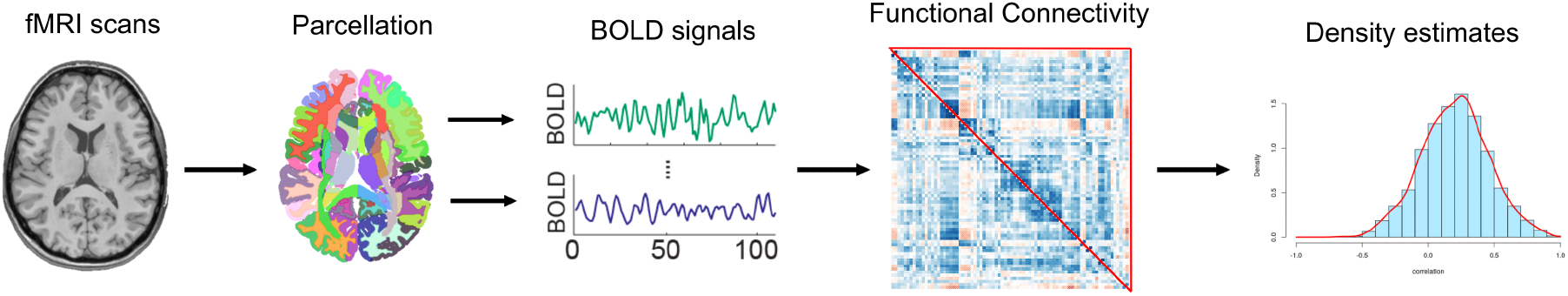
From MRI scan to connectivity density

Our proposal is to use *f_i_* to characterize **C**_*i*_ and subsequently study the relationship between *f_i_* and variables of interest. In the tDCS study, the variable of interest is the treatment status. Since the {*f_i_*} are (infinite dimensional) functional data, we employ functional data analysis tools [22, 23, 24]. Logically, one would model that treatment status predicts connectivity. However, treating complex data as covariates is often more convenient than treating them as the outcomes. Therefore, we adopt the ideas in case-control inverse regression [25, 26], and predict whether a subject is in the treatment arm using the connectivity density and the baseline covariates as predictors. Let *A_i_* denote the treatment assignment with *A_i_* = 1 for tDCS and *A_i_* = 0 for sham, and **X**_*i*_ ∈ ℝ^*q*^ denote the *q*-dimensional covariate vector with the first element one for the intercept. The linear model considered is the following:

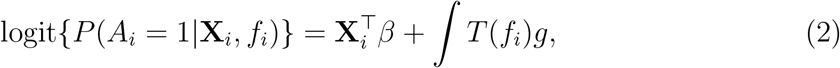

where *T* is a given operator from 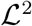 to 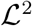 aiming to capture a specific characteristic of the density functions. The function *g* is a coefficient function, and *β* ∈ ℝ^*q*^ is the coefficient vector of the covariates, both to be estimated.

Various choices of *T* and the shape of *g* have different interpretations on the resulting model. For example, setting *T* (*f*) = *f*, the identity function, the linear predictor is ∫*T* (*f_i_*)*g* = *E*[*g*(*Z_i_*)], where *E*[·] is the expectation of a random variable and *Z_i_* is an random variable drawn from *f_i_*. With a sufficiently flexible choice of *g*, mode (2) covers a broad range of possible model fits. However, many of them may not focus on tail behavior, where effects would likely occur. For example, if *g* is a polynomial, the model considers the moments of the density (mean, variance, skewness, etc.) as a predictor. However, it offers no benefit over the direct usage of the moment estimates of the connectivities. Thus, it will not be discussed further, though it does demonstrate a special case of the approach.

As for the choice of *T*, using *T* (*f*) = log(*f*) is similar to the use of the identity function. It loses the expected value interpretation, while instead, performs regression on the space of densities with Aitchison geometry [27]. Thus, it may better detect the influence of the tail behavior on the outcome.

Another choice is the quantile mapping, *T_q_*(*f*) = *F*^−1^, where *F* is the cumulative distribution function associated with the density *f*. With a sufficient number of foci, this approach is approximately equivalent to using the empirical quantiles of the connectivity data as the regressors. Our proposed approach is quite similar to this. However, we further propose to weight the quantiles via density quantile. Specifically, we set *T_ldq_*(*f*) = log ◦ *f* ◦ *F*^−1^ = −log [(*dF*^−1^*/dt*)^*−*1^] where ◦ is the function composition operator. The latter equality is easy to derive by taking derivatives via the chain rule to the identity function, *F* ◦ *F*^−1^. Note that the density quantile *f* ◦ *F*^−1^ can be regarded as a quantile synchronized version of density function, and therefore is more sensitive to the changing tails. And the further logarithm transform maps density quantile to a Hilbert space, which is essential to linear models. This idea has been explored before as a potentially preferable method for utilizing quantiles as regressors. Specifically, it is equivalent to the Hilbert space mapping, suggested by Petersen and Müller [9].

### 2.4. Reversing the predictor/response relationship

It is typical in regression models to consider the hypothetically functionally antecedent variable as a predictor, independent or exogenous variable, rather than an outcome, dependent or endogenous variable. A counterexample is in outcome dependent sampling, such as in retrospective studies. We utilize the same strategy of reversing the typical predictor / response relationship, as is more convenient for modeling with high dimensional and complex quantities (such as brain connectivity) as the predictor. In the tDCS study, we model treatment assignment as the outcome using a logit model with the connectivity density and other covariates as the independent variables. This avoids the need to construct probability distributions on the connectivity densities themselves.

To elaborate, using Bayes’ rule and the fact of a randomization design, *P* (*A_i_* = 1) = *P* (*A_i_* = 0) = 0.5, for any function *g* and transformation *T*, we have:

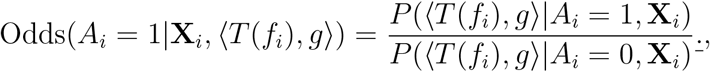

where 〈·, ·〉 is any inner product of two functions. In our application we consider logit models on *P* (*A_i_* = 1|**X**_*i*_, *T* (*f_i_*)), which depends on *f_i_* only though the form 〈*T*(*f_i_*)*, g*〉. As the above relationship shows, our treatment assignment outcome model, *P*(*A_i_*|**X**_*i*_, *T*(*f_i_*)), is consistent with any connectivity outcome model, *P*(〈*T*(*f_i_*)*, g*〉|*A_i_,* **X**_*i*_), where the likelihood ratio comparing treated to controls is approximately log linear with our linear separable density model given in Equation 2.

### 2.5. Estimation of the coefficient function

To estimate the coefficient function, *g* in model (2), we perform a functional principal components analysis (fPCA) [28]. This reduces the dimension of the functional regressor using a set of data-derived basis. In this approach, one calculates the PCA decomposition of the functions, {*f*_*i*_}, using the Karhunen/Loève transformation [29], where the covariance function is smoothed [30] and selects the leading principal components that explain over 99% of the variation as the basis functions. Notice that, the version of fPCA utilized here does not honor possible density implied constraints of the *T*(*f_i_*). Generalized cross validation (GCV) is commonly used to choose the smoothing parameters [for detailed discussion, see Section 4.5.4 of 31]. Confidence bands are derived using a Bayes approach. [32, 33, 24].

### 2.6. Comparison

To illustrate the benefit of conducting a delocalized analysis, a simulation study based on the fMRI data collected in the tDCS study is conducted. We consider an extreme example that demonstrates an example where non-localized brain stimulation decreases statistical power, or even makes it impossible to identify ROI pairs with a significant effect when implementing a localization method. However, using connectivity densities retains the relevant information.

In the simulation, consider a brain connectivity map with 20 regions, *R*_1_ *… R*_20_, a transcranial stimulation that randomly “stimulates” region *R_i_* with equal probability across *i*. After stimulation, the correlations of *R_i_* with all other regions are flipped, with the remaining region pairs unchanged. The mean and variance of the stimualted data are constant across stimulation, mimicking the actual tDCS data. Thus, the stimulation does not impact the first two moments of connectivity and has a very weak localized effect by randomly stimulating different spaces. However, stimulating any region has a consistent impact to connectivity density. This simulation is, of course, an extreme caricature of a non-localized effects in real data.

We sampled 100 pre-stimulation maps from the pre-intervention scans and then simulated 100 post-stimulating maps according to above mechanism. Then, we tested the significance of edgewise testing, the LASSO and density regression, with different transformations. We performed 500 such simulations. For completeness, we also considered these methods when there was no change from before to after stimulation and when the stimulation was localized at a particular region. In the real tDCS data, the density methods are compared with regressing connectivity matrix by comparing edges associated with the estimated connectivity from pairs of foci.

The edgewise regression approach considers the following model:

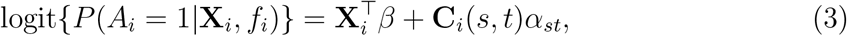

where *s* > *t*. The second competing approach considered was a regression model with high-dimensional predictors:

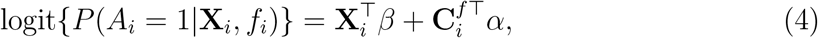

where 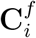 is the vectorization of the upper triangular portion of **C**_*i*_. A LASSO regularization was imposed and high-dimensional inferences were drawn following the procedure introduced by Dezeure et al.[34] We refer to this model as the LASSO model.

## 3. Results

### 3.1. Simulation

Figure 2 shows example connectivity maps and fitted functional regressors from an example simulation, one where stimulation was present and one where it was absent. We report the rate of positive findings for all methods. Results are shown in Table 2. Localization methods do not find significant region pairs in the non-localized simulations. However, the density method detects the stimulation impact on the connectivity densities.

**Figure 2:**
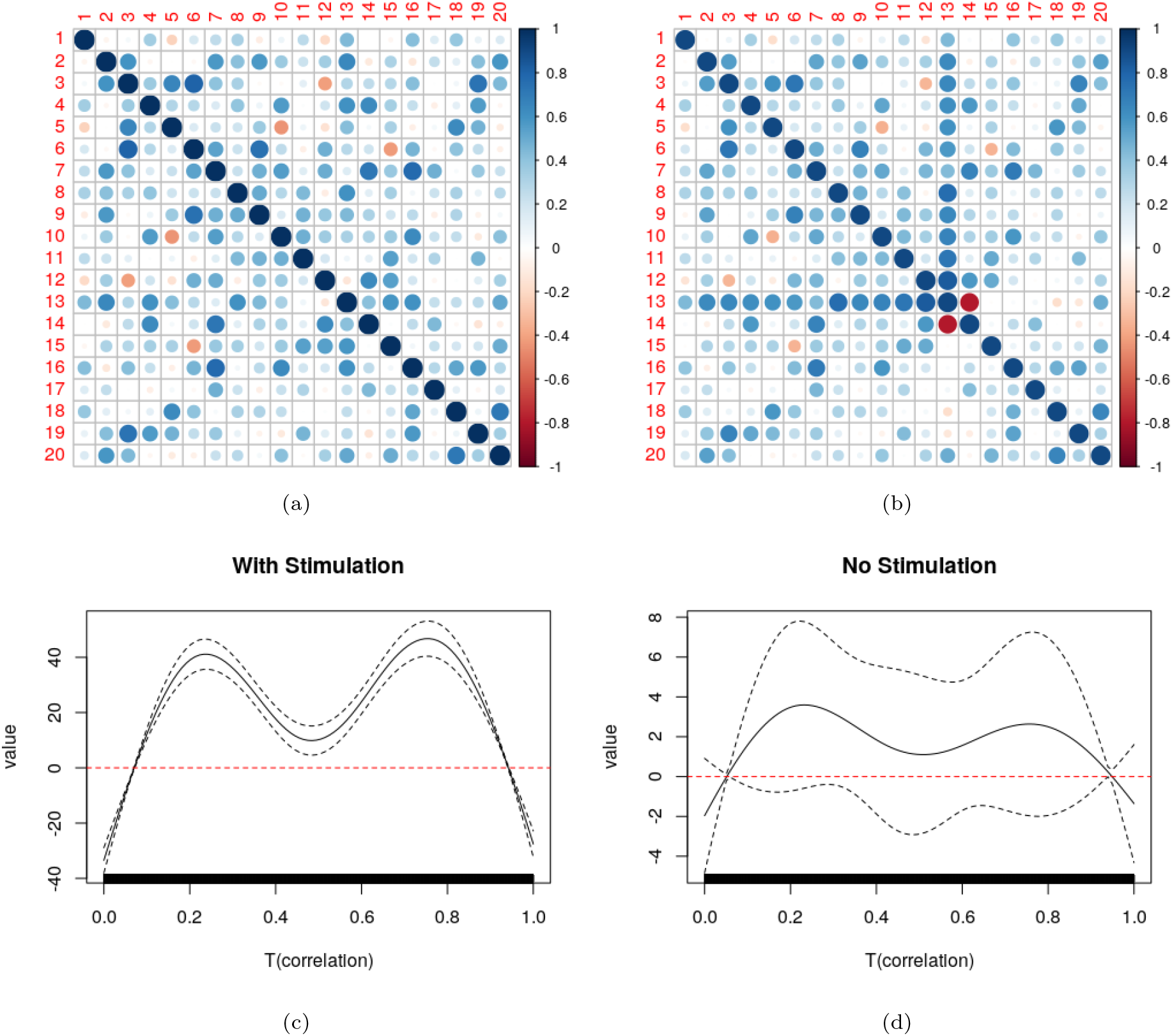
Example simulation results, where (a) and (b) show simulated pre- and post-stimulation connectivity maps. Image (c) shows the regression coefficient function and its 95% confidence interval when there’s stimulation. The regression model uses the log density quantile function. Image (d) shows the same curve when there’s no stimulation.

**Table 2:** This table shows the rate of positive findings over 500 simulations. *T*_0_*, T_l_, T_ldq_* are the identity, logarithm and log density-quantile transformations described in section 2.3. Bonferroni, FDR [35] and BY [36] refer to multiplicity correction procedures. LASSO testing, were implemented by the pacakge hdi [37].

| | Bonferroni | FDR | BY | $T_0$ | $T_l$ | $T_{ldq}$ |
| --- | --- | --- | --- | --- | --- | --- |
| Non-Localized | 0.03 | 0.03 | 0.004 | 1.00 | 1.00 | 1.00 |
| Localized | 1.00 | 1.00 | 1.00 | 1.00 | 1.00 | 1.00 |
| No-Stimulation | 0.01 | 0.01 | 0.00 | 0.05 | 0.07 | 0.04 |

### 3.2. Analysis of the tDCS data using edgewise testing & LASSO

For the tDCS data, we also tested the significance of edgewise regression [models (3), (4)] and a joint model of the upper-triangular component of **C**_*i*_. No foci-pair was identified as significant in either regression model, at Type I error rate levels of 0.05 or 0.1. Of note, previous localization work on related data [38], yields significant findings. However, the total number of regions were restricted, thus dramatically reducing multiplicity concerns. In this analysis, 78 regions were used, resulting in a more stringent correction factor. In addition, a more restrictive inclusion criteria in [38] led to a different study population.

### 3.3. Analysis of the tDCS data using the density regression

In this section, we present the analysis results of the tDCS study using the density regression (Model (2)) with different transformations (*T*). The fitted coefficient function, *g*, and its 95% confidence interval are presented in Figure 3. Functional linear regression was performed using the `refund` R-package with default parameter of smoothed covariance fPCA, which chooses the number of components that explains over 99% of the data variation.

**Figure 3:**
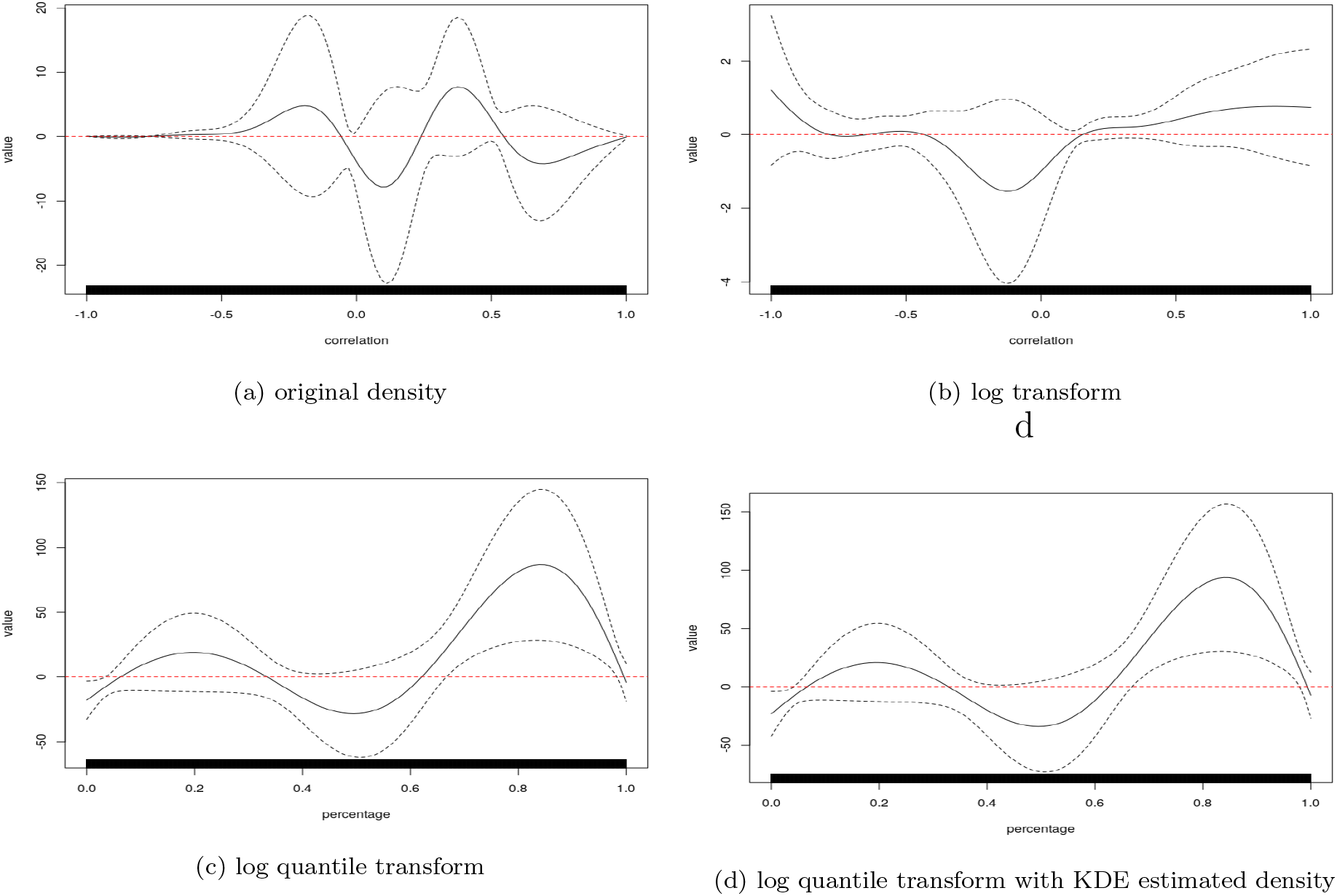
Model results on the tDCS experiment. The black solid line is the fitted coefficient function, *g*, with the black dashed line referencing the associated 95% confidence interval. Densities were estimate from smoothing splines implemented in the **fda** R-package with 19 degrees of freedom for the spline basis. A kernel density estimator (KDE,Figure 3d) is also computed and compared with smoothing spline (Panel 3c) method. Contrasting 3c and 3d shows that the density estimation technique did not impact results.

Regressing on the density after applying the log-density quantile transform yields the highest number of significant signals, which reaches its maximum around the 85^*th*^ percentile. This potentially indicates that stimulation has a consistent tail effect which is more likely to be aligned by quantile, rather than absolute value. Since the estimated coefficient function is significantly non-zero only in the positive tail this suggests that the tDCS group had higher connection densities in the tail than the sham group. That is, connectivity among the most connected regions was higher in the tDCS group.

A likelihood ratio test was performed to compare logistic regression with only baseline variables and our log-quantile model including both the baseline variables and the log density quantile term. The resulting p-value was 0.0052, indicating a statistically significant gain of information from connectivity density at the 0.05 benchmark type I error rate.

### 3.4. Induced Connectivity

Consider the best model using the log density quantile transform, *T_ldq_*. We have

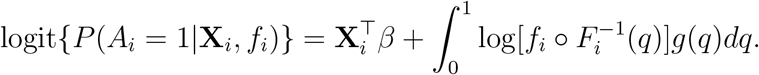

Notice that for the connectivity matrix, **C**_*i*_, we have *F_i_*{**C**_*i*_} ∼ *U*(0, 1), a uniform distribution on [0, 1] via the probability integral transform. Let **Q**_*i*_(*s, t*) = *F*_*i*_{**C**_*i*_(*s, t*)}. Then it follows that:

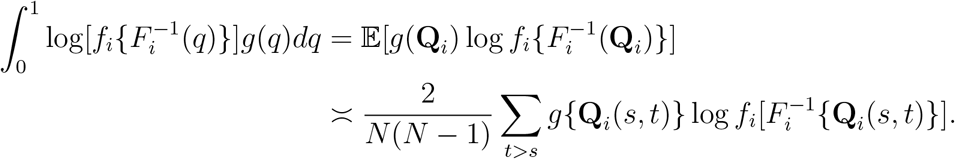

Therefore, for this subject, one can assign 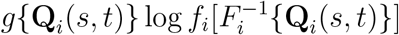 as the effect size for region pair (*s, t*). Averaging this effect across all patients yields an importance metric for every region pair in the model. We call this stimulation induced connectivity, since it describes how influential the correlation of each region pair is in predicting stimulation status. The induced connectivity matrix is shown in Figure 4 together with a summary of effect agreement across subjects, where for each patient, region pairs are selected with top 5% absolute effect size. And the frequency of each region pair being selected is calculated.

**Figure 4:**
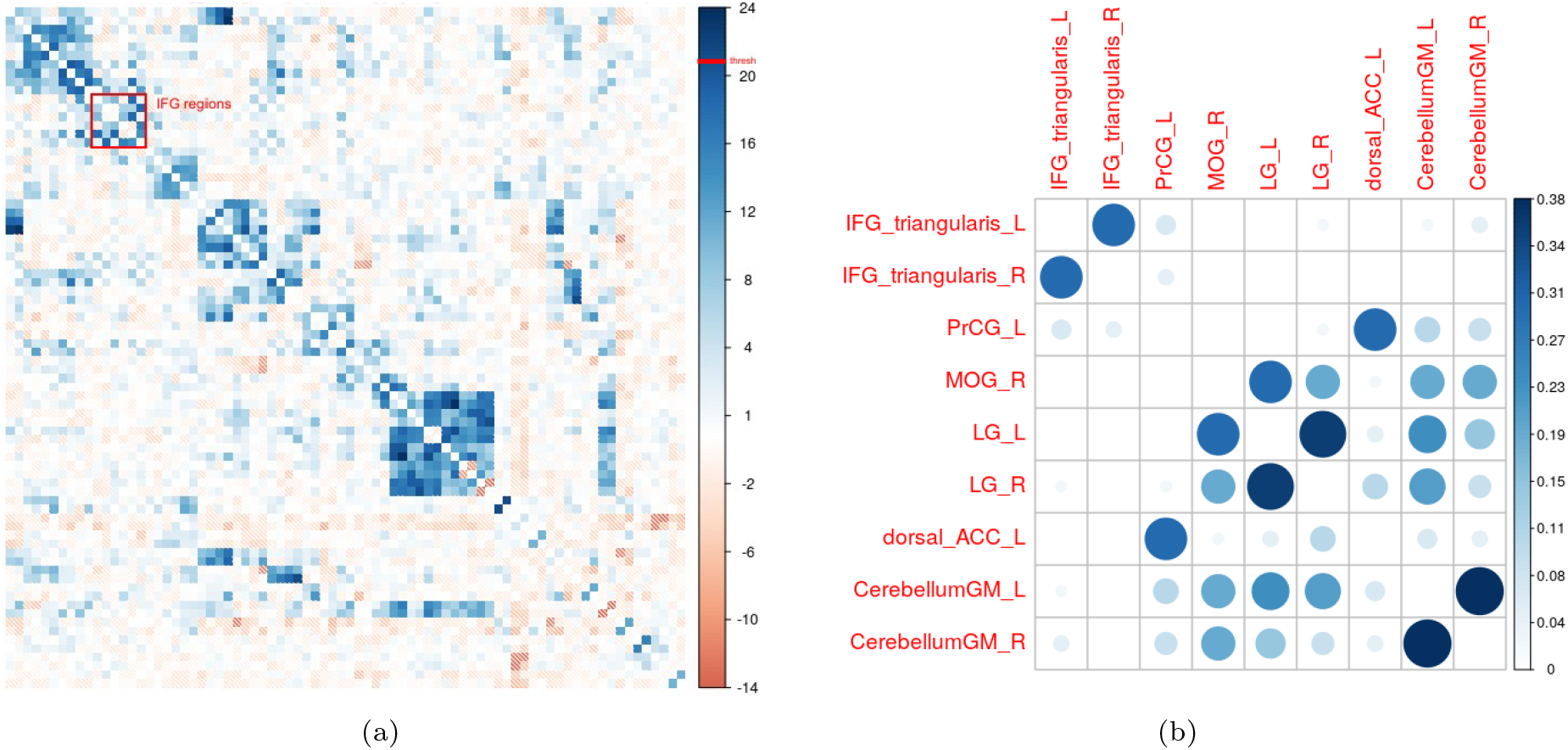
Figure 4a shows the induced connectivity described in section 3.4. IFG regions (which were applied tDCS) are noted in the red box. And figure 4b shows some region pairs with most consistent contribution, measured by the frequency of having top 5% absolute effect size across all patients. Again, IFG regions were the ones applied tDCS

This technique, of course, returns to a discussion of localized effects. However, by investigating this measure one can ascertain the degree of localization consistency across subjects - an impossibility with pure localization analysis.

## 4. Discussion

In this manuscript, a new framework for analysing functional connectivity was proposed. Functional data analysis of log quantile connectivity densities investigates possible non-localized effects associated with subject level variables. A sizable byproduct of this style of analysis is the general elimination of multiplicity considerations. This is of great importance in connectivity analysis, where the number of comparisons grows at a rate of the square of the number of foci being considered. In the data application, we find associations with stimulation and connectivity density. In contrast, edgewise methods fail to find any results purely because of the multiplicity issue. This is partially due to a wide search of all possible region pairs from the parcellation. Of course, one could also reduce multiplicity concerns by restricting attention to regions associated with a priori hypotheses of interest, as was done in [38]. In contrast, investigating connection densities is an omnibus approach that benefits from a reduction in the number of tests over exploratory edge-wise aprpoaches, a robustness to non-localized effects and a robustness to the inclusion of unnecessary foci. These benefits come at the expense of the loss of power and interpretability over analyese considring only a small set of tightly specified edge-wise hypotheses.

An interesting direction to pursue with connectivity density methods is to consider robustness to spatial registration [39]. The connectivity density should be largely invariant to registration. In contrast, localization methods heavily rely on both accurate registration and accurate biological functional localization across subjects. Therefore, density regression could be performed after affine registration typically done prior to the more time consuming non-linear registration.

We used functional data analysis to relate connection densities to outcomes. Functional data analysis tools [23] have grown to be quite flexible. Thus, density regression approaches can be relatively easily generalized to handle different settings, such as any typical statistical outcome model and longitudinal data. Also, density estimates may naturally make adjustments for missing data, in the form of missing foci, since the density can remain the same in some contexts. This has potential broad implications for the study of stroke and other diseases with abnormal brain pathology. Localization methods are not available if the region of interest is damaged or missing. In contrast, density based methods are easy to apply.

Statistically, we assumed independence between subjects and relied on the randomization to invert the predictor / response relationship using logit models. This borrows techniques from case referent sampling from epidemiology dating back to the seminal work of Cornfield [40, 41]. Independence between subjects was used for inference. We also used density estimates for connection densities, techniques that implicitly require sampling assumptions for theoretical convergence. However, we contend that connectivity densities are intrinsically of interest, and therefore no appeals to super-population inference and sampling assumptions are needed for estimation. This is analogous to spatial group ICA, where productive estimates are obtained via independence assumptions on voxels over space, without a true sampling or super-population model for inference [42]. An interesting future direction of research would investigate dependencies between foci correlations.

Our recommended approach uses log quantile densities as the functional predictor, rather than the density, distribution function or quantile function directly [9]. This approach has convenient theoretical properties, but also the practical benefit of focusing attention on tail behavior, where effects are most likely to be seen. Utilizing the quantile density also creates robustness to irrelevant foci pairs being included in the analysis.

Our simulations and data results focus on settings that highlight the benefits of an omnibus density regression approach. In the simulations, we investigated a non-localized caricature of typical effects. Similarly, in our data analysis, we performed no filtering of regions prior to analysis (thus magnifying multiple comparison concerns). It was shown in the simulation, that functional density regression approaches can find real non-localized effects, whereas, as expected, edgewise methods do not find any. It should be emphasized that the performance of the density regression approach is invariant to the distribution of effects across subjects, whereas edgewise approaches become viable as the degree of localization increases.

In addition, the flexibility of the approach finds tail effects in the real data, even though there are a great deal of irrelevant connections (i.e. unnecessarily included region pairs) being studied. Edgewise and other regression approaches are highly sensitive to unnecessary null connections being included in the analysis. A benefit of the data being considered is the likely existence of an effect related to the stimulation. However, we emphasize that a single omnibus approach does not represent a full analysis of the data. We recommend this approach as a global analysis to be performed prior to edgewise or other localization methods. This mirrors the classic ANOVA (analysis of variance) approach of performing an overall F test before investigating pairs of explanatory factor levels. It would most useful in exploratory model building where foci selection is not restrictive. In cases of tightly coupled statistical hypotheses involving relatively few regions or foci, density regression would not be needed or particularly helpful.

This methodology raise many avenues for future research. For example, one the idea of non-localized effects in dynamic connectivity [43] via stochastic processes of connectivity densities (by time). In addition, there are multiple alternatives for densities estimated from correlation of each region pair for contralateral regions. Here, it should be acknowledged that there is strong homotopic correlations from symmetric regions. One should then deal with multivariate densities estimated from pairs of correlations. This same logic could be applied to geographically close regions and for instances with longitudinal scans. The connectivity density of spectral information [44], like leading principal component scores, should also be studied to potentially extract relevant brain graph properties.

Finally, there’s the role that connecvity density methods could play in fMRI analysis of subjects with missing brain tissue, such as studies of stroke or surgical interventions. Connectivity density methods may be resilient to the missing data impact of differential brain structure in a way that localization methods are not. In fact, it is interesting to conjecture what localization methods even mean in these settings where a subset of subjects are missing areas of localization. In contrast, density methods may provide a more robust and well defined methodology. It is worthy of note that components of graph methodology [45, 45, 46] often considers summary metrics that do not require or assume localization. Density regression can be considered a subset of weighted graph metric analysis.

## 5. Acknowledgments

We would like to thank our participants and referring physicians for their dedication, helpful comments and interest in our study.

## References

[1] K. J. Friston, Functional and effective connectivity: a review, Brain connectivity 1 (2011) 13–36.

[2] J. S. Damoiseaux, M. D. Greicius, Greater than the sum of its parts: a review of studies combining structural connectivity and resting-state functional connectivity, Brain Structure and Function 213 (2009) 525–533.

[3] A. M. Bastos, J.-M. Schoffelen A tutorial review of functional connectivity analysis methods and their interpretational pitfalls, Frontiers in systems neuroscience 9 (2016) 175.

[4] D. Koutra, J. T. Vogelstein, C. Faloutsos Deltacon: A principled massive-graph similarity function, in: Proceedings of the 2013 SIAM International Conference on Data Mining, SIAM, 2013, pp. 162–170.

[5] J. T. Vogelstein, W. G. Roncal, R. J. Vogelstein, C. E. Priebe, Graph classification using signal-subgraphs: Applications in statistical connectomics, IEEE transactions on pattern analysis and machine intelligence 35 (2012) 1539–1551.

[6] P. Broca Remarques sur le siège de la facultédu langage articulé, suivies d’une observation d’aphémie (perte de la parole), Bulletin et Memoires de la Societe anatomique de Paris 6 (1861) 330–357.

[7] C. Wernicke Der aphasische Symptomencomplex: eine psychologische Studie auf anatomischer Basis, Cohn., 1874.

[8] S. Finger C. R. Almli, Brain damage and neuroplasticity: mechanisms of recovery or development?, Brain Research Reviews 10 (1985) 177–186.

[9] A. Petersen H.-G. Müller, et al., Functional data analysis for density functions by transformation to a hilbert space, The Annals of Statistics 44 (2016) 183–218.

[10] D. Scheinost J. Benjamin C. Lacadie B. Vohr K. C. Schneider, L. R. Ment, X. Papademetris R. T. Constable, The intrinsic connectivity distribution: a novel contrast measure reflecting voxel level functional connectivity, Neuroimage 62 (2012) 1510–1519.

[11] M. A. Nitsche, L. G. Cohen, E. M. Wassermann, A. Priori N. Lang A. Antal W. Paulus F. Hummel P. S. Boggio, F. Fregni et al., Transcranial direct current stimulation: state of the art 2008, Brain stimulation 1 (2008) 206–223.

[12] M. L. Gorno-Tempini, A. E. Hillis, S. Weintraub A. Kertesz M. Mendez S. F. Cappa, J. M. Ogar, J. Rohrer S. Black B. F. Boeve, et al., Classification of primary progressive aphasia and its variants, Neurology 76 (2011) 1006–1014.

[13] D. S. Knopman, J. H. Kramer, B. F. Boeve, R. J. Caselli, N. R. Graff-Radford, M. F. Mendez, B. L. Miller, N. Mercaldo Development of methodology for conducting clinical trials in frontotemporal lobar degeneration, Brain 131 (2008) 2957–2968.

[14] S. Mori D. Wu C. Ceritoglu Y. Li A. Kolasny M. A. Vaillant, A. V. Faria, K. Oishi M. I. Miller, Mricloud: delivering high-throughput mri neuroinformatics as cloud-based software as a service, Computing in Science & Engineering 18 (2016) 21–35.

[15] C. Ceritoglu X. Tang M. Chow D. Hadjiabadi D. Shah T. Brown M. H. Burhanullah, H. Trinh J. Hsu K. A. Ament, et al., Computational analysis of lddmm for brain mapping, Frontiers in neuroscience 7 (2013) 151.

[16] M. I. Miller, M. F. Beg, C. Ceritoglu C. Stark Increasing the power of functional maps of the medial temporal lobe by using large deformation diffeomorphic metric mapping, Proceedings of the National Academy of Sciences 102 (2005) 9685–9690.

[17] X. Tang K. Oishi A. V. Faria, A. E. Hillis, M. S. Albert, S. Mori M. I. Miller, Bayesian parameter estimation and segmentation in the multi-atlas random orbit model, PloS one 8 (2013) e65591.

[18] A. V. Faria, S. E. Joel, Y. Zhang K. Oishi P. C. van Zjil, M. I. Miller, J. J. Pekar, S. Mori Atlas-based analysis of resting-state functional connectivity: Evaluation for reproducibility and multi-modal anatomy–function correlation studies, Neuroimage 61 (2012) 613–621.

[19] Y. Behzadi K. Restom J. Liau T. T. Liu, A component based noise correction method (compcor) for bold and perfusion based fmri, Neuroimage 37 (2007) 90–101.

[20] C. Gu C. Qiu Smoothing spline density estimation: Theory, The Annals of Statistics (1993) 217–234.

[21] B. W. Silverman, Density estimation for statistics and data analysis, volume 26, CRC press, 1986.

[22] J. O. Ramsay, Functional data analysis, Encyclopedia of Statistical Sciences 4 (2004).

[23] J. O. Ramsay, B. W. Silverman, Applied functional data analysis: methods and case studies, Springer, 2007.

[24] M. W. McLean, G. Hooker A.-M. Staicu F. Scheipl D. Ruppert Functional generalized additive models, Journal of Computational and Graphical Statistics 23 (2014) 249–269.

[25] R. L. Prentice, R. Pyke Logistic disease incidence models and case-control studies, Biometrika 66 (1979) 403–411.

[26] K. J. Rothman, S. Greenland T. L. Lash, Case–control studies, Encyclopedia of Quantitative Risk Analysis and Assessment 1 (2008).

[27] J. J. Egozcue, J. L. Düaz-Barrero, V. Pawlowsky-Glahn, Hilbert space of probability density functions based on aitchison geometry, Acta Mathematica Sinica 22 (2006) 1175–1182.

[28] P. T. Reiss, R. T. Ogden, Functional principal component regression and functional partial least squares, Journal of the American Statistical Association 102 (2007) 984–996.

[29] R. G. Ghanem, P. D. Spanos, Stochastic finite elements: a spectral approach, Courier Corporation, 2003.

[30] C.-Z. Di C. M. Crainiceanu, B. S. Caffo, N. M. Punjabi, Multilevel functional principal component analysis, The annals of applied statistics 3 (2009) 458.

[31] S. N. Wood, Stable and efficient multiple smoothing parameter estimation for generalized additive models, Journal of the American Statistical Association 99 (2004) 673–686.

[32] G. Wahba Bayesian “confidence intervals” for the cross-validated smoothing spline, Journal of the Royal Statistical Society: Series B (Methodological) 45 (1983) 133–150.

[33] D. Nychka Bayesian confidence intervals for smoothing splines, Journal of the American Statistical Association 83 (1988) 1134–1143.

[34] R. Dezeure P. Bühlmann, L. Meier N. Meinshausen High-dimensional inference: Confidence intervals, p-values and r-software hdi, Statistical science (2015) 533–558.

[35] Y. Benjamini Y. Hochberg Controlling the false discovery rate: a practical and powerful approach to multiple testing, Journal of the Royal statistical society: series B (Methodological) 57 (1995) 289–300.

[36] Y. Benjamini D. Yekutieli et al., The control of the false discovery rate in multiple testing under dependency, The annals of statistics 29 (2001) 1165–1188.

[37] N. Meinshausen L. Meier P. Bühlmann, P-values for high-dimensional regression, Journal of the American Statistical Association 104 (2009) 1671–1681.

[38] B. N. Ficek, Z. Wang Y. Zhao K. T. Webster, J. E. Desmond, A. E. Hillis, C. Frangakis A. V. Faria, B. Caffo K. Tsapkini The effect of tdcs on functional connectivity in primary progressive aphasia, NeuroImage: Clinical 19 (2018) 703–715.

[39] F. P. Oliveira, J. M. R. Tavares, Medical image registration: a review, Computer methods in biomechanics and biomedical engineering 17 (2014) 73–93.

[40] N. E. Breslow, Statistics in epidemiology: the case-control study, Journal of the American Statistical Association 91 (1996) 14–28.

[41] S. W. Greenhouse, Jerome cornfield’s contributions to epidemiology, Biometrics (1982) 33–45.

[42] V. D. Calhoun, T. Adali G. D. Pearlson, J. J. Pekar, A method for making group inferences from functional mri data using independent component analysis, Human brain mapping 14 (2001) 140–151.

[43] R. M. Hutchison, T. Womelsdorf E. A. Allen, P. A. Bandettini, V. D. Calhoun, M. Corbetta S. Della Penna, J. H. Duyn, G. H. Glover, J. Gonzalez-Castillo, et al., Dynamic functional connectivity: promise, issues, and interpretations, Neuroimage 80 (2013) 360–378.

[44] W. de Haan, W. M. van der Flier, H. Wang P. F. Van Mieghem, P. Scheltens C. J. Stam, Disruption of functional brain networks in alzheimer’s disease: what can we learn from graph spectral analysis of resting-state magnetoencephalography?, Brain connectivity 2 (2012) 45–55.

[45] O. Sporns Networks of the Brain, MIT press, 2010.

[46] E. Bullmore O. Sporns Complex brain networks: graph theoretical analysis of structural and functional systems, Nature reviews neuroscience 10 (2009) 186–198.

